# Targeted quantification of phosphorylation sites identifies STRIPAK-dependent phosphorylation of the Hippo pathway-related kinase SmKIN3

**DOI:** 10.1101/2020.12.17.423311

**Authors:** Valentina Stein, Bernhard Blank-Landeshammer, Ramona Märker, Albert Sickmann, Ulrich Kück

## Abstract

We showed recently that the germinal <u>c</u>entre <u>k</u>inase III (GCKIII) SmKIN3 from the fungus *Sordaria macrospora* is involved in sexual development and hyphal septation. Our recent extensive global proteome and phosphoproteome analysis revealed that SmKIN3 is a target of the <u>str</u>iatin <u>i</u>nteracting <u>p</u>hosphatase <u>a</u>nd <u>k</u>inase (STRIPAK) multi-subunit complex. Here, using protein samples from wild type and three STRIPAK mutants, we applied absolute quantification by <u>p</u>arallel <u>r</u>eaction <u>m</u>onitoring (PRM) to analyze phosphorylation site occupancy in SmKIN3 and other <u>s</u>eptation <u>i</u>nitiation <u>n</u>etwork (SIN) components, such as CDC7 and DBF2, as well as BUD4, acting downstream of SIN. For SmKIN3, we show that phosphorylation of S668 and S686 is decreased in mutants lacking distinct subunits of STRIPAK, while a third phosphorylation site, S589, was not affected. We constructed SmKIN3 mutants carrying phospho-mimetic and phospho-deficient codons for phosphorylation sites S589, S668 and S686. Investigation of hyphae in a ΔSmKin3 strain complemented by the S668 and S686 mutants showed a hyper-septation phenotype, which was absent in the wild type, the ΔSmKin3 strain complemented with wild type gene, or the mutant S589. Furthermore, localization studies with SmKIN3 phosphorylation variants and STRIPAK mutants showed that SmKIN3 preferentially localizes at the terminal septa, which is distinctly different from the wild type strains. We conclude that STRIPAK-dependent phosphorylation of SmKIN3 has an impact on controlled septum formation and on the time-dependent localization of SmKIN3 on septa at the hyphal tip. Thus, STRIPAK seems to regulate SmKIN3, as well as DBF2 and BUD4 phosphorylation, affecting septum formation.

## Introduction

The STRIPAK multi-subunit complex functions as a macromolecular assembly communicating through physical interactions with other conserved signaling protein complexes to constitute larger dynamic protein networks. STRIPAK is involved in a broad variety of developmental processes in higher and lower eukaryotes. For example, proliferation of several mammalian cancer cells is correlated with dysfunctional STRIPAK subunits (1–3), and in fungal microorganisms, the lack of STRIPAK results in sexual infertility, defects in hyphal fusion, and impaired pathogenicity or symbiotic interactions (4).

We are interested in identifying putative phosphorylation and dephosphorylation targets of STRIPAK in the filamentous fungus *S. macrospora*, a filamentous ascomycete closely related to *Neurospora crassa (5)*. Techniques to globally quantify the proteome and phosphoproteome, such as <u>l</u>abel-<u>f</u>ree <u>q</u>uantification (LFQ) and label-based approaches (<u>i</u>sobaric <u>T</u>ags for <u>R</u>elative and <u>A</u>bsolute <u>Q</u>uantitation, iTRAQ, <u>t</u>andem <u>m</u>ass <u>t</u>ag, TMT), are indispensable for large-scale detection of changes in phosphorylation of peptides and the identification of potential molecular targets of kinases and phosphatases (6). We recently performed extensive isobaric tagging for relative and absolute quantification-based proteomic and phosphoproteomic analyses to identify potential targets of STRIPAK in *S. macrospora*. The proteome and phosphoproteome of the wild type and three different STRIPAK deletion mutants revealed a total of 4,193 proteins and 2,489 phosphoproteins in all strains, where 1,727 proteins were present in both proteomes. Among these, we identified 781 phosphoproteins that showed differential phosphorylation in all mutants compared to the wild type (7). However, the functional role of the posttranslational protein modifications was only characterized in a few cases (7,8).

Certain inherent limitations with these shotgun methods, such as ratio compression and under sampling, can only be overcome by the complementary use of targeted mass spectrometry (MS) approaches. These can be employed as a means to validate a subset of the results obtained by shotgun experiments. While targeted MS is widely used for accurate protein quantification, the high variability, increased experimental effort, and need for validation has limited the implementation of targeted approaches in phosphoproteomics analysis (9). A hallmark study showed that 25% of differentially regulated phosphopetides were attributed to alterations at the protein level (10). Thus, to accurately determine the phosphorylation ratio of a given site, targeted quantification of phosphorylation sites is highly appropriate to quantify both the corresponding phosphorylated and non-phosphorylated peptides in order to obtain site-specific phosphorylation ratios (11,12).

Among the 781 regulated proteins in *S. macrospora* mentioned above (7), we found the GCKIII SmKIN3, which however was absent in the global proteome, and thus did not permit quantitative measurement of phosphorylation. Therefore, SmKIN3 phosphorylation was determined by applying absolute quantification by synthesis of <u>s</u>table <u>i</u>sotope-labeled <u>s</u>tandard (SIS) peptides combined with PRM, a method which to the best of our knowledge has not yet been applied to a fungal organism.

SmKIN3 is involved in septation of hyphae and associated with the highly conserved SIN (13). The SIN complex, homologous to Hippo signaling in animals, comprises a STE-kinase, a GCK and a <u>n</u>uclear <u>D</u>BF2-<u>r</u>elated (NDR) kinase (14). For example, in *Neurospora crassa* the STE-kinase CDC-7 phosphorylates GCK SID-1, the homolog of SmKIN3, and activates DBF-2 (15). The function of SIN is essential for septation and cytokinesis, as demonstrated by SIN-deletion strains (13,15,16). BUD4, an anillin-related protein, acts further downstream and specifies SIN-regulated septum formation (17,18).

Here, we show that STRIPAK-directed dephosphorylation of SmKIN3 has a significant impact on proper hyphal septation and septal localization. To the best of our knowledge, this is the first report about STRIPAK-dependent phosphorylation analyzed by targeted quantification of phosphorylation sites, and will have an impact on understanding the function of mammalian homologs.

## Results

### Absolute quantification of phosphorylation site occupancy by PRM of germinal centre kinase SmKIN3

Previously, we showed that the GCKIII SmKIN3 is associated with STRIPAK and regulates fungal development (13). Sequence comparison of the primary amino acid sequence showed a homology of 92.43% with the corresponding sequence from *N. crassa*; but similarity with other homologues from ascomycetes is low. Using the <u>e</u>ukaryotic <u>l</u>ength <u>m</u>otif (ELM) database (19), we found a kinase domain in the amino terminus, several LATS kinase recognition motifs, and a binding motif for <u>f</u>ork<u>h</u>ead<u>a</u>ssociated (FHA) domains. The COILS sequence analysis program (http://www.ch.embnet.org/software/COILS_form.html), revealed two predicted putative coiled-coil domains located next to each other in a region between amino acids 688 and 788. At the C-terminal end of SmKIN3 (amino acids 805-811), we detected a conserved sequence motif, previously called T-motif (Fig. 1*A*). Such T-motifs have only been found so far in a small family of related fungal kinases (20). Database research revealed that a motif occurs for example in SmKIN3, its homologue Sid1p from *Schizosaccharomyces pombe* and SID-1 from *N. crassa*.

**Figure 1:**
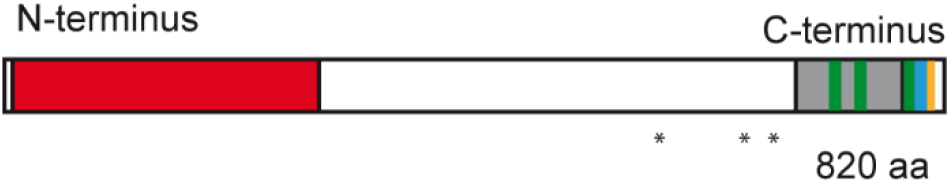
Linear structure of SmKIN3 and identified protein domains. Depicted are a serine/threonine kinase domain (aa 10-279, red), large tumor suppressor kinase 1 (LATS1) kinase phosphorylation motifs (aa 721-727; 740-746; 794-800, green), a coiled-coil domain (aa 688-788, grey), a T-motif (aa 801-806, blue), and a phospho-threonine motif, binding a subset of FHA domains (aa 805-811,yellow). Asterisks below indicate phosphorylation sites S589, S668, and S686.

Recently, we determined the phosphoproteome of *S. macrospora* wild type and STRIPAK mutant strains (7,8), and detected three phosphorylation sites S589, S668 and S686 in the SmKIN3 sequence, which are conserved between *S. macrospora* and *N. crassa* (Fig. 1*B*). Due to its low abundance in the overall proteome, we were unable to quantify the STRIPAK-dependent phosphorylation of SmKIN3 (7,8).

Here, we set out to determine the impact of STRIPAK on SmKIN3 phosphorylation, by applying targeted quantification of phosphorylation site occupancy using PRM. As described above, SmKIN3, together with CDC7 and DBF2 constitute the SIN complex, which is associated with the downstream landmark protein BUD4. Therefore, we included all these components in our PRM analysis.

A targeted bipartite TiO_2_-LC-PRM-based workflow was established to quantify the site-specific phosphorylation states of putative STRIPAK targets. First, 70 phosphorylated peptides and their corresponding non-phosphorylated counterparts were selected and SIS were synthesized. The median <u>c</u>oefficient of <u>v</u>ariation (CV) of the biological replicates was calculated as 16.3% for all non-phosphorylated, and 16.1% for all phosphorylated peptides. To further prioritize our approach, we selected 15 pairs of peptides, representing phospho-sites from proteins belonging to the SIN signaling pathway. Dilution series were prepared to verify a linear response and determine the <u>l</u>imit <u>o</u>f <u>d</u>etection (LOD) of each SIS peptide, as described in the Methods section. LODs ranged from 1.7 amol to 19.3 fmol per injection for phosphorylated SIS peptides, and from 1.43 amol to 32.3 fmol per injection for their non-phosphorylated counterparts. The average <u>l</u>ower <u>l</u>imit <u>o</u>f <u>q</u>uantification (LLOQ) was 375 amol or 135 amol on-column for phosphorylated or non-phosphorylated peptides. The median CV throughout all dilution steps was calculated as 6.7% for phosphorylated, and 5.0% for non-phosphorylated peptides. These results are summarized in Supplementary data file S1.

*S. macrospora* wild type and the three STRIPAK deletion strains Δ*pro11,* Δ*pro22* and Δ*pp2Ac1* were grown in triplicates prior to being subjected to lysis and protein extraction (Fig. 2). After tryptic digestion and quality control measurements, protein amounts were normalized and aliquots were spiked with either phosphorylated SIS peptides or their non-phosphorylated counterparts. The phosphopeptide aliquots were subjected to TiO_2_-based enrichment followed by LC-PRM measurement, while the fraction spiked with the non-phosphorylated peptides was measured directly. The targeted PRM measurements of SIS and endogenous peptides allowed the simultaneous quantification of both the phosphorylated and non-phosphorylated peptide counterparts, and thus enabled us to calculate the phosphorylation site occupancy.

**Figure 2:**
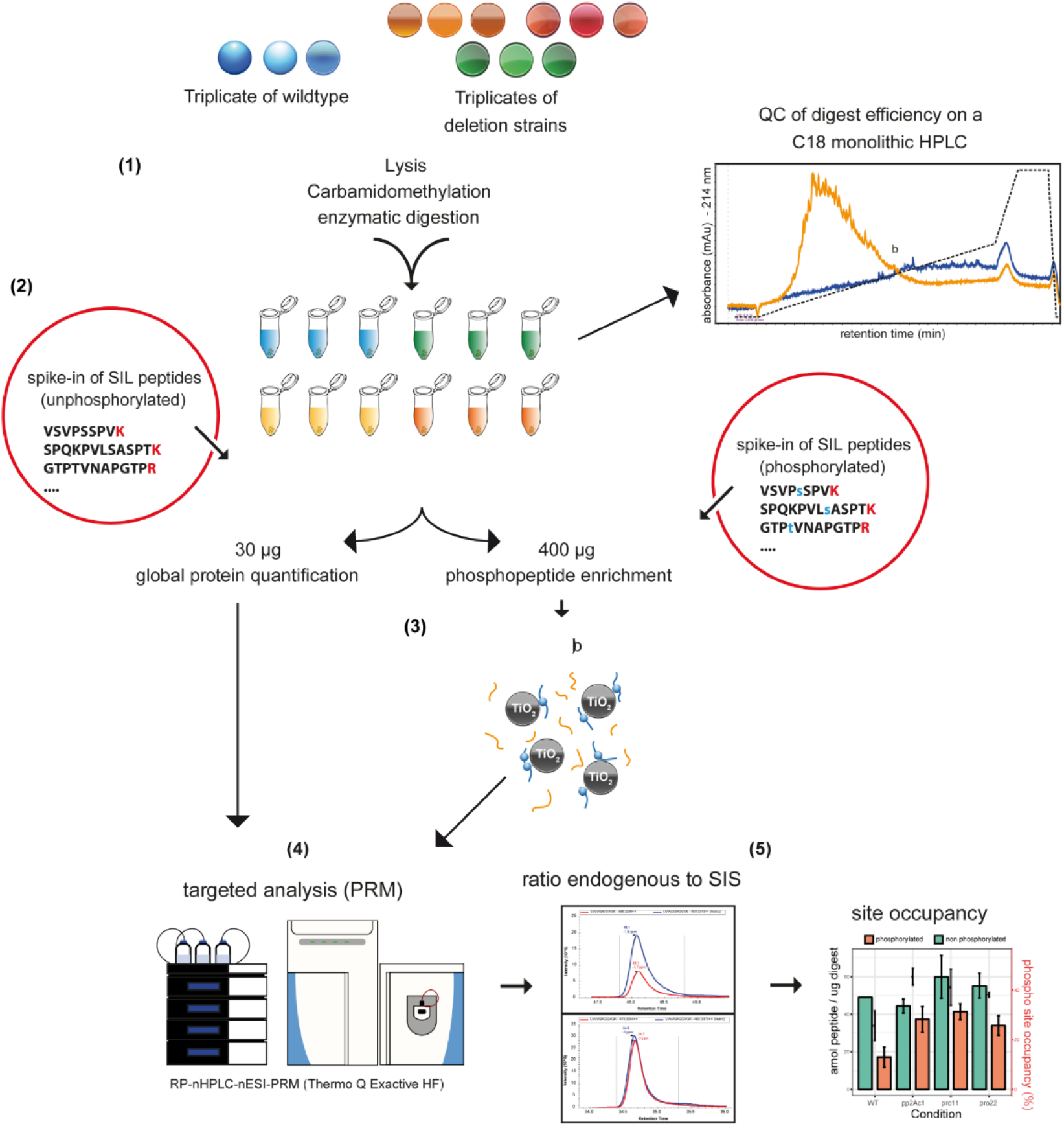
Targeted quantification of phosphorylation site occupancy. Graphical representation of the proteomics workflow including (1) extraction and digestion of proteins from *S. macrospora* strains grown in triplicates, (2) spike-in of phosphorylated and unphosphorylated SIS peptides, (3) TiO_2_-based enrichment of phosphorylated peptides, (4) targeted analysis using a RP-nHPLC-nESI-PRM setup, and (5) data analysis and calculation of phosphorylation site occupancy.

Using the proteomics workflow described above, we were able to distinguish three different outcomes. First, we determined the site occupancy for 37 phosphorylation sites on 4 proteins, where the phosphorylated and non-phosphorylated peptides were determined. Included is the phosphorylation site S686 from SmKIN3. For DBF2, we detected four phosphorylation sites of which three were quantified. Phosphorylation site S104 in DBF2 was significantly decreased when STRIPAK is non-functional, indicating that this site is STRIPAK dependent. Phosphorylation of sites S89 and S502 was not differentially regulated in STRIPAK deletion mutants (Dataset S2, Fig. S1-S3). We found five phosphorylation sites in BUD4 (T381, S367, S369, S742, S1373); all were dephosphorylated in STRIPAK deletion mutants (Dataset S2, Figs. S4-S7). Secondly, nine phosphorylation sites were detected on five proteins; however, the non-phosphorylated peptide was not detectable. The phosphorylation site occupancy for these peptides was indirectly calculated based on the concentration of the remaining peptide pairs of the respective protein. Among these we found two sites (S668, S589) in the SmKIN3 protein. Finally, as a third option, we found seven phosphorylation sites on five proteins, where only the phosphorylated peptides were detected and no site occupancy value could be calculated (Dataset S2). However, changes in the abundance of selected phosphorylated peptides could still be used for quantification, since the corresponding proteins were stably expressed in wild type and STRIPAK deletion mutants in previous experiments. An example is the CDC7 protein, which was recently detected in a phosphoproteomic analysis (7)

Overall, 19 phosphorylation sites showed significantly different site occupancy between the wild type and at least one of the STRPIAK mutant strains (student’s t-test p-value < 0.05; the results are summarized in Supplementary data file S2). This quantification of site occupancy helped us to prioritize our functional analyses of phosphorylation sites from SmKIN3, the main objective of this investigation. It was also the source for constructing a mechanistic model of phosphorylation-dependent protein regulation. The number of peptides where both phosphorylation sites S668 and S686 are dephosphorylated in STRIPAK deletion mutants is higher than in wild type, i.e. peptides containing phosphorylated site S668 and S686 are less abundant in STRIPAK deletion mutants than in the wild type (Fig. 3). In contrast, we detected only a slightly higher number of peptides with phosphorylation at site S589 (Fig. S8).

**Figure 3:**
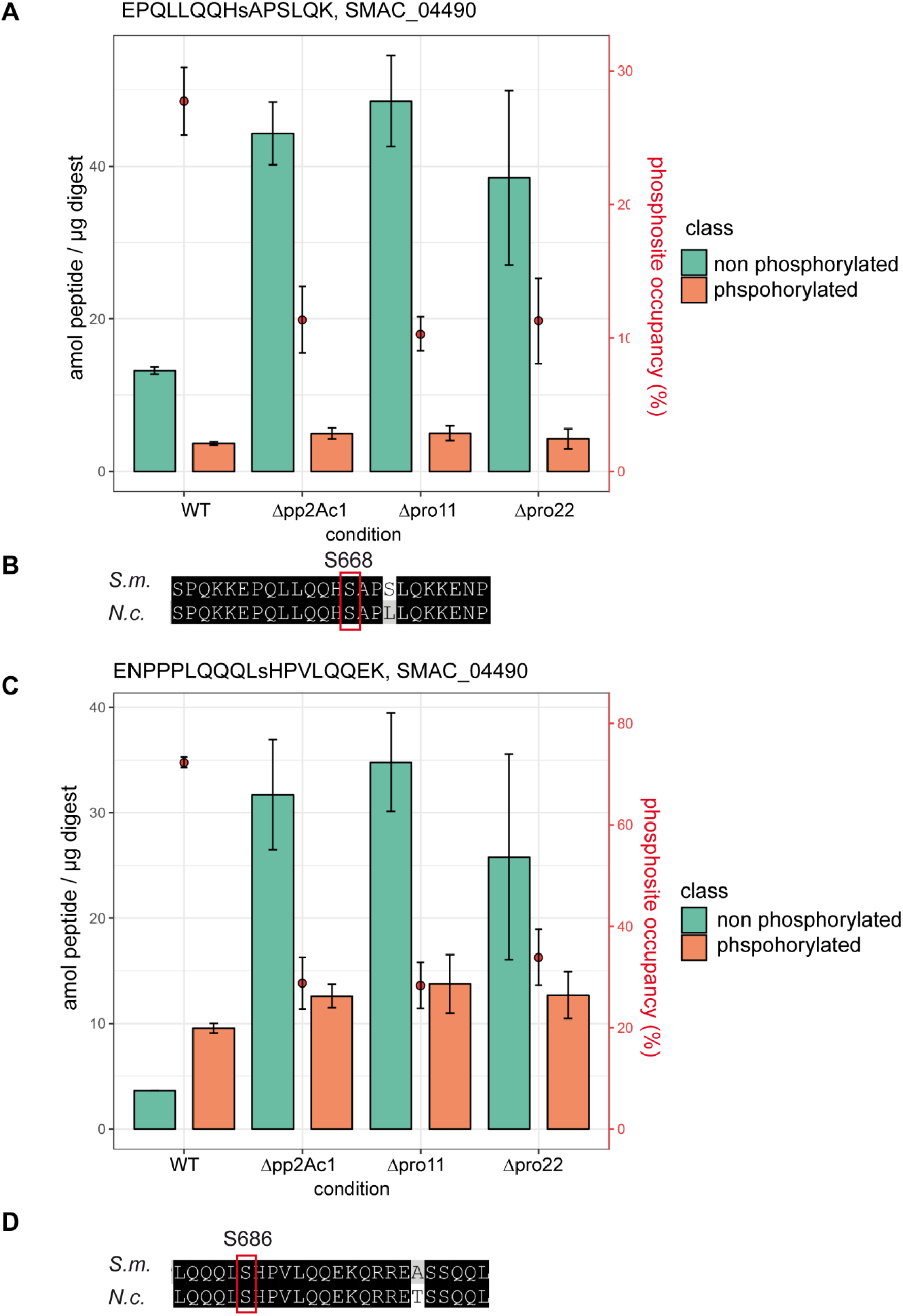
Quantitative analysis of the non-phosphorylated and phosphorylated peptides containing phosphorylation site S668 and S686 of SmKIN3 in three different STIPAK deletion mutants. (A, C) The y-axis on the left gives the amount of the peptides EPQLLQQHsAPSLQK or ENPPPLQQQLHsHPVLQQEK in amol as non-phosphorylated (green) and phosphorylated variants (orange). The y-axis on the right shows the quantity of phosphosite occupancy in percent, plotted as red dots in the bar chart. Error bars indicate standard deviation. Lower case letters in the peptides indicate the phosphosite. (B, D) Alignment of sequences of SmKIN3 from *S. macrospora (S.m.),* and SID-1 from *Neurospora crassa (N.c.).*

### Phospho-mimetic and phospho-deficient SmKIN3 mutants display a hyper-septation phenotype

Our quantitative analysis of the phosphosite occupancy in SmKIN3 indicated that SmKIN3 is a target of STRIPAK. To analyze the function of phosphorylated SmKIN3, we generated six different phospho-deficient plus phospho-mimetic mutants, by subjecting the triplets encoding S589, S668 and S686 to *in vitro* mutagenesis, substituting the serine triplets by either alanine (no phosphorylation) or glutamic acid (mimics phosphorylation) triplets, as described in the Material and Methods section (Fig. S9). Phosphorylation of S589 is apparently STRIPAK independent (Fig. S8), while S668 and S686 seem to depend on STRIPAK (Fig. 3).

For functional analysis of SmKIN3 and its variants, we used a previously described *Smkin3* deletion strain for complementation analysis (13). Recombinant plasmids (Table 2) encoding SmKIN3-GFP phospho-variants were transformed into ΔSmKin3, and primary transformants were used to isolate homokaryotic ascospores. The expression of SmKIN3-GFP wild type and phospho-variants was verified by Western blot analysis indicating the expression of a 118 kDa protein (Fig. S10). When the wild type tagged *Smkin3* gene was used for complementation, we obtained fully fertile strains with wild-type mycelial growth, indicating that GFP-labeled SmKIN3 is fully functional. We then investigated three homokaryotic ascospore isolates of each phospho-mimetic strain, S589E, S668E, S686E, and phospho-deficient strain, S589A, S668A and S686A. Fluorescence microscopy showed that phospho-mutated SmKIN3 still localizes to septa and close to nuclei and thus resembles the wild type situation (Fig. S11-S17). However, we found that both S668A and S668E showed an increased number of septa in hyphal branches, as detected by GFP fluorescence of the SmKIN3-phospho variants. The identity of septa was further confirmed by <u>C</u>alco<u>f</u>luor <u>W</u>hite (CFW) staining. We investigated at least 200 single hyphal branches for each strain. In these cases, 600 hyphal branches were counted for each recombinant strain (Fig. 4*A-C*, Dataset S3). We counted all septa within 20 μm behind hyphal branches. Double, triple, quadruple and quintuple septa were considered when they were located within a distance of maximum 12 μm. These are referred to as hyper-septation phenotype.

**Fig 4:**
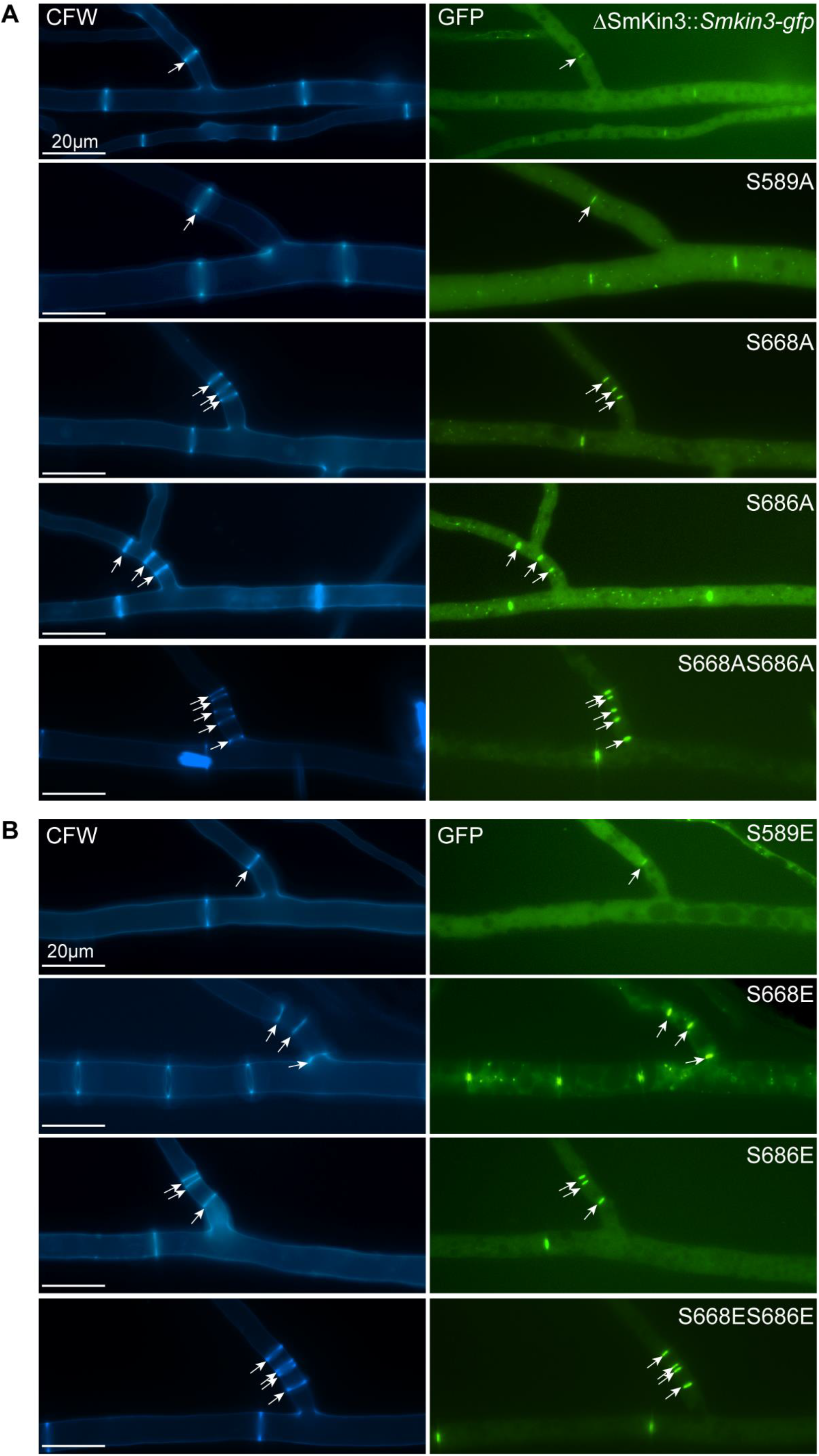
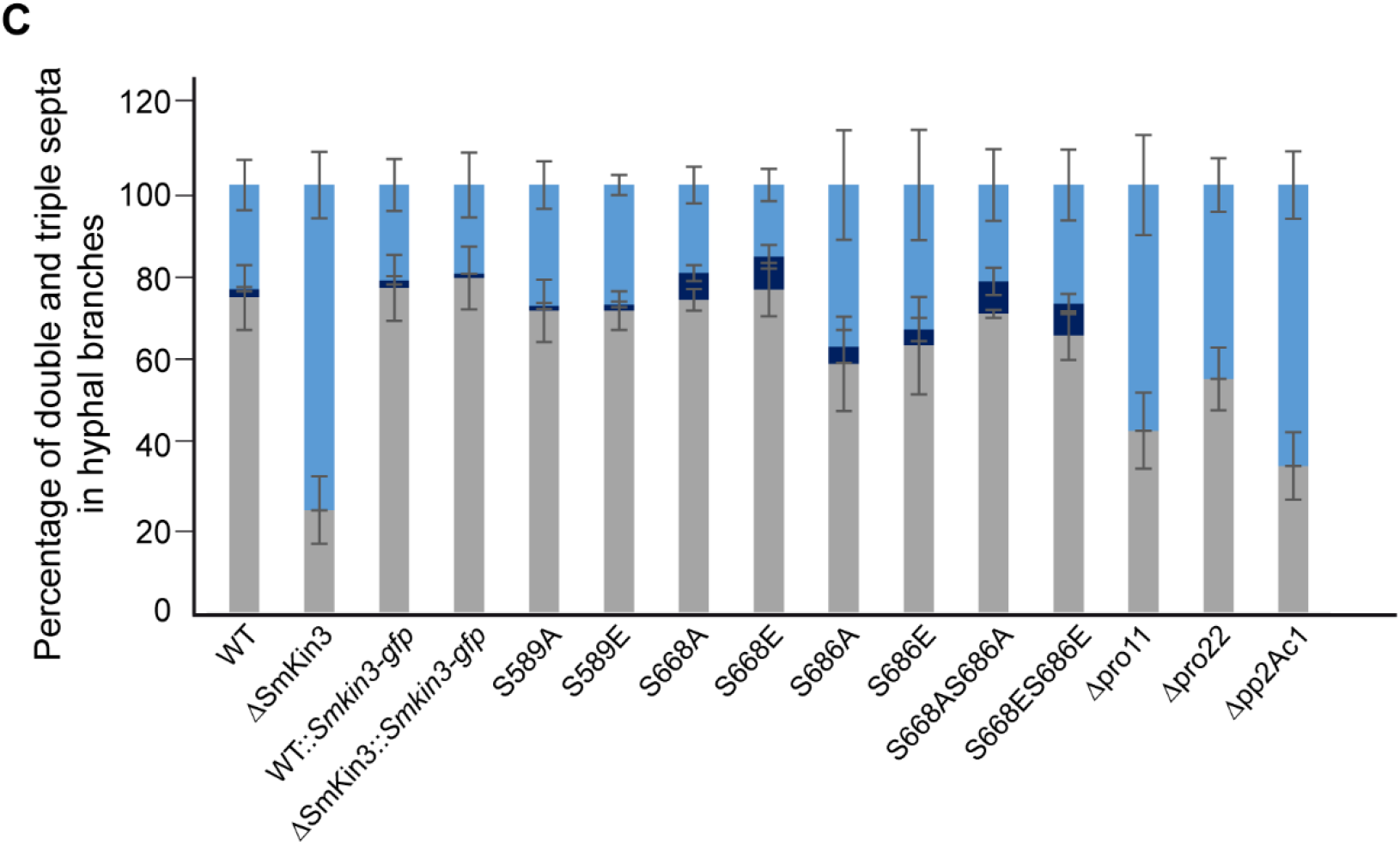
Hyper-septation phenotype in wild type and phospho-mutants. (A) Fluorescence microscopic investigation of septation at hyphal branches. Mycelia were stained with CFW, and SmKIN3 is labeled with GFP. (B) Quantitative investigation of hyper-septation. All values are given in percent. We investigated at least 100 hyphal branches for each STRIAPK mutant strain. In the case of recombinant strains, three different single ascospore isolates were investigated to exclude side effects from random integration of recombinant DNA into genomic DNA. In these cases, 600 hyphal branches were counted for each recombinant strain. As a control, 600 hyphae from ΔSmKin3 complemented with wild type, and ΔSmKin3 strains were investigated in 3 technical replicates. Percentage of hyperseptation (2-5 septa) are given in dark blue, no septa are indicated in light blue. Error bars indicate standard deviations.

In the case of the hyper-septation, the value for the *S. macrospora* wild type was 1.84% ± 0.50. The hyper-septation value for the complemented strains and phospho-mutants were as follows: WT::*Smkin3-gfp* 1.00% ± 0.69, ΔSmKin3::*Smkin3-gfp* 3.08% ± 2.90, phospho-mutants S589A 1.04% ± 0.64, S589E 1.44% ± 0.63, S668A 5.80% ± 1.69, S668E 6.96% ± 2.53, S686A 4.00% ± 3.80, S686E 3.41% ± 2.50.

Obviously, mutations of S668 and S686 have an effect on hyphal septation. To verify their effect further, we constructed phospho-mimetic (S668ES686E) and phospho-deficient (S668AS686A) double mutants and analyzed derived ascospore isolates. The strains were fully fertile and the phospho-mutated SmKIN3 proteins are still found at septa and at nuclei in young hyphae (Fig. S18-19). We determined the corresponding values for hyper-septations: S668AS686A 7.00% ± 2.99, S668ES686E 6.84% ± 2.14. Interestingly, we detected triple, quadruple and quintuple septa only in the phosphomutants S668 and S686, and the double mutants S668S686, but they were lacking in wild type, strains complemented with the wild type gene, or phospho-mutants S589A and S589E.

From our quantitative results we conclude that the phospho-mimetic and phospho-deficient SmKIN3 mutations at S686 and S668 are responsible for the hyper-septation phenotype, while phosphorylation site S589 has no effect on septum formation. Interestingly, the investigated double mutants resemble mostly to mutants S668. To analyze the impact of STRIPAK on septum formation, we examined three STRIPAK deletion strains lacking either the regulatory subunit of the STRIPAK phosphatase (*Δpro11*), the catalytic subunit of the STRIPAK phosphatase (*Δpp2Ac1)*, or the STRIP1/2 homologue (*Δpro22*). We found no hyper-septation in *Δpro22* and *Δpp2Ac1* and only a very low number of double septa in *Δpro11* (0.8% ± 1.1) (Fig. 4*C*, Fig. S20. We conclude that septum formation is reduced in STRIPAK mutants, and thus is probably positively regulated by STRIPAK.

### Localization of SmKIN3 at septa is dependent on phosphorylation and an intact STRIPAK complex

Next, we investigated the effect of phosphorylation of SmKIN3 on its cellular localization at septa. For fluorescence and differential interference microscopy we used all eight phosphorylation variants as well as the wild type and two deletion strains lacking genes for STRIPAK subunits. We selected the four terminal septa at the hyphal tip for our detailed localization analysis. We counted the first terminal septa at hyphal tips, where GFP fluorescence indicated SmKIN3 localization (representative images in Fig. 5). As an example, more than 50% of wild type septa showed fluorescence at the 3^rd^ terminal septum (Table 1). Similar values were obtained for the ΔSmKin3 strain complemented with the wild type *Smkin3* gene. However, ΔSmKin3 strains, carrying the mutated codon S589 were different; most of the localization was observed at the 2^nd^ and 3^rd^ septum. A significantly different result was observed with all mutated codons S668, S686 as well as with the double mutants S668S686. Here, we found preferential localization at the first or second septum (Fig. S23).

**Fig. 5:**
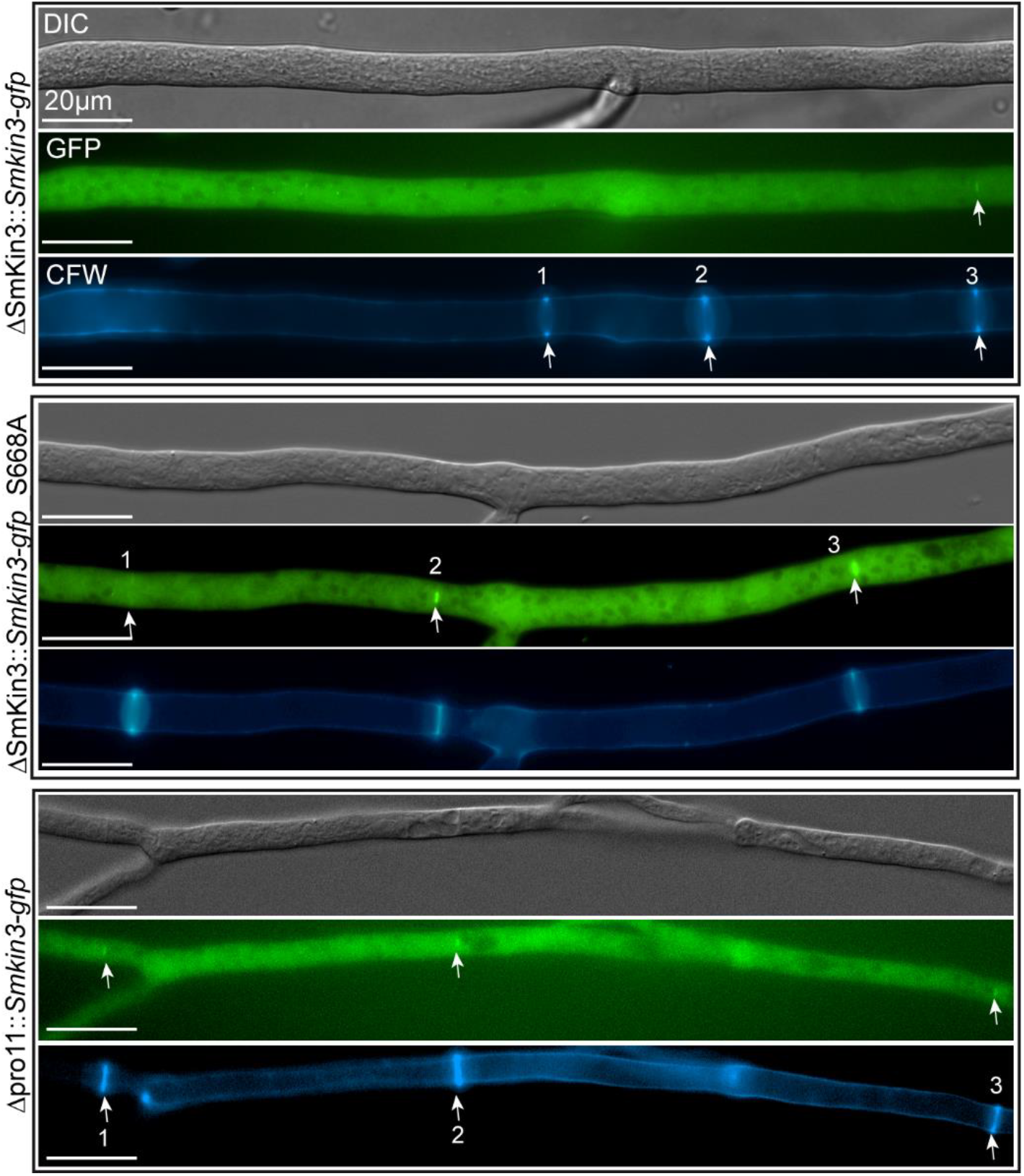
STRIPAK-dependent localization of SmKIN3-GFP at hyphal septa. The strains shown here are indicated on the left. The first three septa from the hyphal tip (on the left) were numbered. Arrowheads label green fluorescing SmKIN3 (GFP), or CFW stained septa.

**Table 1:**
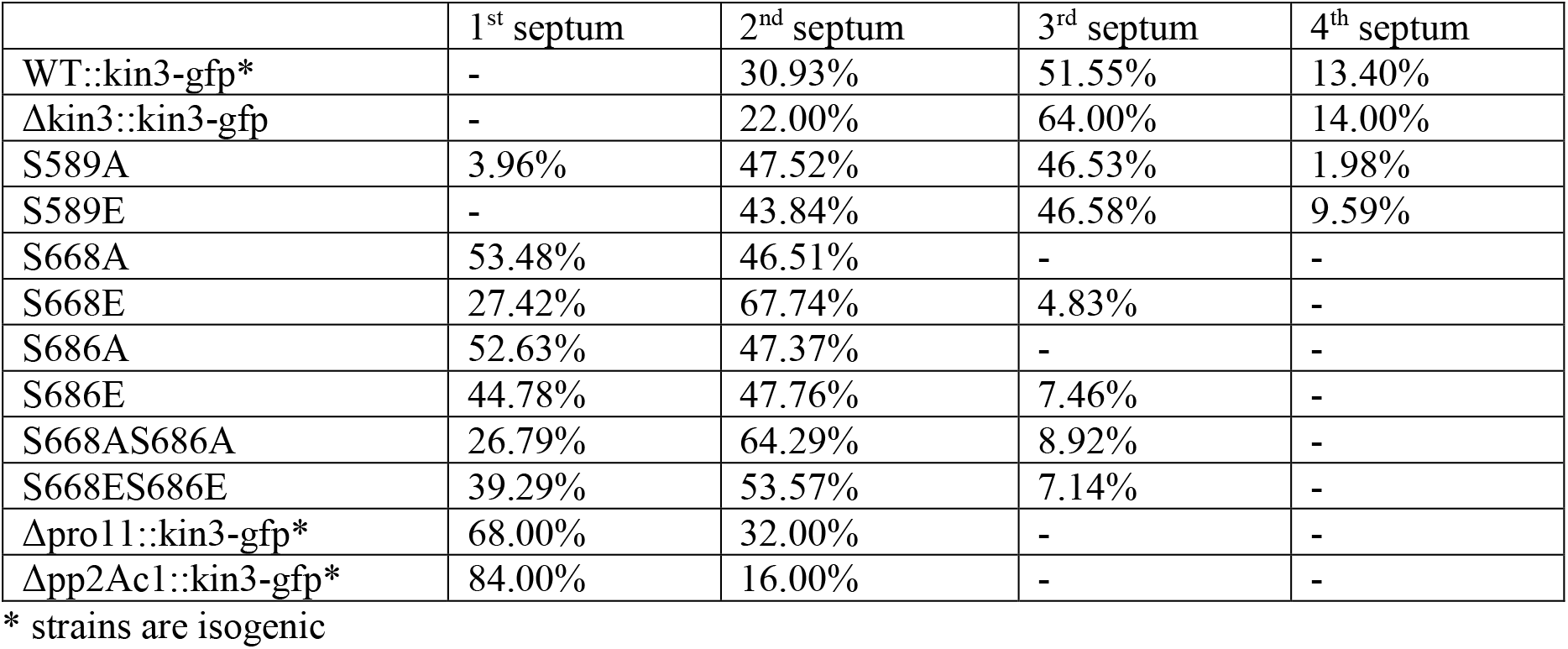
Localization of SmKIN3, and its phosphorylation variants in wild type and STRIPAK deletion strains. In wild type, SmKIN3 is preferentially seen at the third terminal septum, while in the three variants S668, S686 and S668S686, and both STRIPAK mutants *Δpp2Ac1* and *Δpro11*, localization is observed mostly at the first or second septum. N≥ 50 per strain. Localization of SmKIN3 in the complemented ΔSmKin3 strain and in both phospho-variants of S589 resembles the wild type.

|  | 1<sup>st</sup> septum | 2<sup>nd</sup> septum | 3<sup>rd</sup> septum | 4<sup>th</sup> septum |
| --- | --- | --- | --- | --- |
| WT::kin3-gfp* | - | 30.93% | 51.55% | 13.40% |
| $\Delta kin3::kin3-gfp$ | - | 22.00% | 64.00% | 14.00% |
| S589A | 3.96% | 47.52% | 46.53% | 1.98% |
| S589E | - | 43.84% | 46.58% | 9.59% |
| S668A | 53.48% | 46.51% | - | - |
| S668E | 27.42% | 67.74% | 4.83% | - |
| S686A | 52.63% | 47.37% | - | - |
| S686E | 44.78% | 47.76% | 7.46% | - |
| S668AS686A | 26.79% | 64.29% | 8.92% | - |
| S668ES686E | 39.29% | 53.57% | 7.14% | - |
| $\Delta pro11::kin3-gfp^*$ | 68.00% | 32.00% | - | - |
| $\Delta pp2Ac1::kin3-gfp^*$ | 84.00% | 16.00% | - | - |

\* strains are isogenic

To further investigate the role of STRIPAK, we analyzed the localization of SmKIN3 in two STRIPAK deletion mutants, *Δpro11* and *Δpp2Ac1*. In both deletion strains, SmKIN3 mainly localized at the first or second septum, similar to the phospho-variants of S668 and S686, and unlike in the wild type strain. This suggests that temporally controlled phosphorylation on sites S668 and S686 is required for the localization of SmKIN3 to early septation sites. The microscopic study was further extended by demonstrating that SmKIN3 also localized to septa in mature hyphae (Figs. S11-S17, S22-S24).

## Discussion

Here, we provide analytical molecular evidence that phosphorylation of SmKIN3 at distinct sites is STRIPAK dependent. Tightly regulated phosphorylation of SmKIN3 is important for its function. Not only did we discover that deregulation of SmKIN3 phosphorylation leads to a hyper-septation phenotype, but also that temporal phosphorylation of SmKIN3 regulates its septal localization.

We accurately quantified 53 phosphorylated peptides, and for 37 of those directly determined the site occupancy of the phosphorylation sites by quantifying the corresponding non-phosphorylated peptide. To date, accurate targeted quantification and determination of phosphorylation site occupancy has only been performed in several studies, and these are mostly limited to just a handful of sites (11,21–24), or only used crude SIS peptides (25). Since our targeted phosphorylated peptides were found at concentrations lower than 30 amol per μg of protein lysate, to the best of our knowledge this result confirms that we have used a most sensitive assay to determine phosphorylation site occupancy in a non-human model organism.

### STRIPAK regulates SmKIN3 phosphorylation indirectly

In mammalian cells, Hippo comprises two kinases, GCK MST1/2 and NDR kinase LATS1/2 (26,27). Recently, it was shown that STRIPAK integrates upstream signals to control the activity of the SmKIN3 homolog MST1/2 for initiating Hippo signaling. Deletion of STRIP1/2, a homolog of *S. macrospora* PRO22, results in upregulation of MST1/2, which led to the conclusion that STRIPAK regulates MST1/2 (26,28,29).

In fungi, the Hippo homologous kinase cascade SIN comprises three kinases. For example, in *N. crassa* the Ste20 kinase CDC-7 acts upstream of GCK SID-1 and NDRK DBF-2 (30). SID-1, the middle component of SIN, is the homolog of SmKIN3 and MST1/2. Our PRM analysis revealed that the phosphosite occupancy, i.e. phosphorylation in two out of three sites in SmKIN3 is decreased in STRIPAK deletion mutants. This finding is intriguing, since STRIPAK is a dephosphorylating complex. However, comparable results in fission yeast showed that SIN is negatively regulated by the <u>S</u>IN <u>i</u>nhibitory <u>P</u>P2A (SIP) complex, the STRIPAK homolog (31). SIP dephosphorylates the upstream Ste20 kinase Cdc7p, which leads to assembly of SIN. Cdc7p itself phosphorylates SID1p, the homolog of SmKIN3. Dysfunction of SIP prevents assembly of SIN and thus abolishes phosphorylation of SID1p (31). Sid1p, SID-1 and SmKIN3 not only resemble each other in their posttranslational modifications, but also in their functions. Similar to Sid1p in *S. pombe*, SmKIN3 and SID-1 in *N. crassa* are required for proper septum formation (15,31).

### The SIN component SmKIN3 is a positive regulator of septum formation

In filamentous ascomycetes, such as *S. macrospora*, hyphae are compartmentalized by the formation of septa, which are assembled at an actomyosin-based <u>c</u>ortical <u>r</u>ing (CR), followed by CR constriction. The CR protein complex includes structural proteins, molecular motors, and signaling enzymes (17). Since SmKIN3 localizes at the centre of the hyphal septum around the pore, SmKIN3 may be involved in the function of CR signaling enzymes to activate or inhibit targets via posttranslational modifications, such as phosphorylation. In particular, the presence of SmKIN3 at mature septa suggests additional functions besides septum formation, as was hypothesized for BUD4 (32).

Our PRM approach provides evidence that SmKIN3 is a dephosphorylation target of STRIPAK. The analysis of phospho-mimetic and phospho-deficient SmKIN3 expressing strains revealed a hyper-septation phenotype. Since phosphorylation at distinct SmKIN3 sites is regulated by STRIPAK, we hypothesize that the phosphorylation status of S668 and S686 is important for the temporal and spatial fine tuning of the septation process, and the correct formation of septa.

In *A. nidulans,* SEPH, the homolog of CDC7, has been identified as a central component for the initiation of septation prior to actin ring formation. SEPH is the upstream kinase of SEPL, which is the homolog of SmKIN3, followed by SIDB (homolog of DBF2) (Bruno et al. 2001; Kim et al. 2006). SIN and its downstream effectors are involved in forming the CR, which is responsible for initiating the formation of septa. To specify the location for septum formation, axial landmark proteins such as BUD3 and BUD4 are needed, which recruit septin AspB to the CR (17). Downregulation of SepH abolishes septation, whereas hyper-activation results in the formation of multiple septa (Bruno et al. 2001). This indicates that SEPH acts as a positive regulator of SIN, which triggers cytokinesis in *A. nidulans* (33). Comparable phenotypes were observed in *S. macrospora,* where the loss of *Smkin3* or mutation in the ATP-binding site results in the reduction of septa (13). Phosphorylation mutants of SmKIN3, as reported here, show a hyper-septation phenotype that was similarly observed in *A. nidulans* mutants with hyper-activation of SEPH. Our results are consistent with findings for *A. nidulans* and *N. crassa*, which showed that homologs of CDC7 and SmKIN3 act in the same pathway as positive regulators of SIN.

### The phosphorylation state of SmKIN3 affects its localization

Here, we demonstrate that SmKIN3’s septal localization is altered in mutated phospho-deficient strains, and STRIPAK deletion mutants, indicating that this process is also mediated by STRIPAK. Furthermore, the STRIPAK-dependent phosphorylation state of SmKIN3 affects its affinity for septal proteins and thus its localization. These findings are consistent with results obtained in fission yeast. There, it was shown that SIN and STRIPAK affect each other by phosphorylation and dephosphorylation (31). SIN phosphorylates GEFs and GTPases, which in turn phosphorylate the formin SepA. The phosphorylation of SepA is important for the assembly of actin to form the CR. The NDR kinase Sid2p is necessary to phosphorylate the septin Cdc12 and other proteins to form the CR (34). Such phosphorylation at the CR must be tightly regulated to form the final septum.

To summarize our findings, we have designed a schematic mechanistic model, as depicted in Fig. 6. The SIN complex acts downstream of STRIPAK, which dephosphorylates CDC7 directly (31). However, STRIPAK acts indirectly on phosphorylation sites S668 and S686 of SmKIN3, while phosphorylation of S589 is likely regulated by an unknown phosphatase and kinase. Furthermore, STRIPAK also indirectly phosphorylates S104 in DBF2, while S89 and S502 are not STRIPAK dependent. Finally, from our PRM analysis, we propose that phosphorylation of all sites from BUD4 depends on STRIPAK.

**Figure 6:**
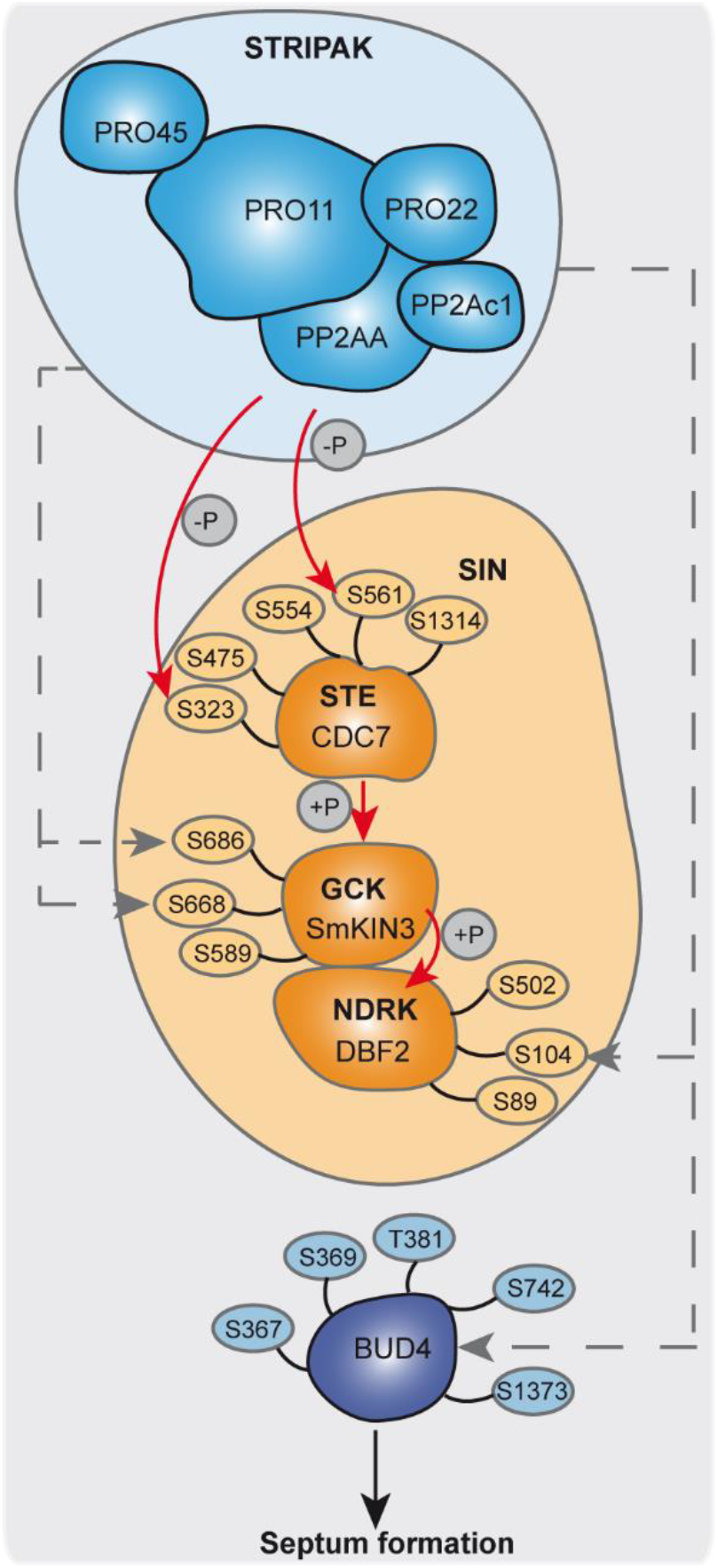
Mechanistic model of STRIPAK-dependent phosphorylation of SmKIN3, as part of the SIN complex. Relevant phosphorylation sites in SIN components and the related BUD4 protein are indicated around the proteins. STRIPAK subunits are colored in blue, SIN components are colored in orange. Grey dashed arrows indicate STRIPAK regulated phosphorylation. Red arrows indicate direct phosphorylation. This model is based on data from this study, and from a recent publication (7)

We hypothesize further that deletion of STRIPAK subunits results in higher phosphorylation of CDC7, which prevents assembly of SIN (Singh et al. 2011). This results in a lower level of phosphorylated SmKIN3, which consequently is unable to phosphorylate the downstream kinase DBF2. Subsequently, DBF2 does not phosphorylate the anillin-like protein BUD4 to form the CR, leading to lower septation. Indeed, our quantitative phosphorylation data provide evidence that four phosphorylation sites of BUD4 and the phosphorylation site S104 of DBF2 are dephosphorylated in STRIPAK deletion mutants compared to wild type. The phosphorylation site S502 of DBF2, which is the homolog to the phosphorylation site S499 in *N. crassa,* is not regulated by STRIPAK (Supplementary data S2). This is consistent with results obtained in *A. nidulans*. There, localization of septin AspB, which is the homolog of CDC12 in *S. macrospora*, is dependent on the formin SepA (SMAC_04496), the SIN kinase SepH (CDC7), and on its own phosphorylation state.

Dephosphorylation of conserved threonine 68 in SepH was shown to be critical for timing of septation and localization (35,36). In *N. crassa,* phosphorylation analysis revealed that phosphorylation of DBF-2 is stimulated by SID-1. Interestingly, phospho-deficient and phospho-mimetic mutants of DBF-2 phosphorylation site S499 in *N. crassa* are nonfunctional *in vivo* and reduce the kinase activity of DBF-2 *in vitro* (15). However, our data indicate that S499 (S502 in *S. macrospora*) seems not to be regulated by STRIPAK, but is essential in septum formation.

The interaction between components of the SIN cascade is decreased when their dephosphorylation is diminished due to a non-functional STRIPAK complex. Moreover, the lack of SIN phosphorylation prevents the recruitment of SIN components to septal pore proteins at the hyphal tip (37). Thus, the interaction of SmKIN3 with SIN and STRIPAK has to be strongly regulated to form septa in a wild type-like manner, as suggested by our localization experiments with wild type and mutant strains.

SmKIN3 homologs are conserved from yeast to humans, and SIN-like pathways have been mentioned in this discussion. We propose that the mechanism of SmKIN3 function described in this study may be applicable to that of homologous networks in other organisms.

## Experimental procedures

### Strains and growth conditions

Electro-competent *E. coli* cells XL1 Blue MRF’ were used for cloning and propagation of recombinant plasmids (38) under standard laboratory conditions (39). All *S. macrospora* strains used in this study are listed in Table S1 and were grown under standard conditions (40,41). For analysis of distribution, SmKIN3 strains were grown for 24 h on solid <u>b</u>io<u>m</u>alt-cornmeal <u>m</u>edium (BMM)-coated glass. Isogenic and homokaryotic strains were generated by genetic crossing and ascospore isolation (40). To obtain phospho-mutants, the mutated plasmids (Table S2) were transformed into a Δ*Smkin3* strain. Phospho-mutations in the generated strains were verified by PCR analysis and DNA sequencing (Eurofins Genomics; Ebersberg, Germany). The number of ectopically integrated recombinant DNA fragments was verified by Southern hybridization analysis (Fig. S24).

### *In vitro* recombinant techniques

Plasmids used in this study are listed in Table S2. pIG1783-Sm*kin3-gfp* was created by restricting the plasmid pIG1783 with *Nco*I. *Smkin3* was amplified via PCR with overhangs at the 5’ and 3’ ends containing recognition sites for *Nco*I. Ligation was performed with T4 DNA ligase. For phospho-mimetic and phospho-deficient strains, plasmid pIG1783-Sm*kin3-gfp* was used for Q5 mutagenesis (NEB biolabs). With specific primers (Table S3), we generated eight plasmids, containing phospho-mimetic and phospho-deficient mutations (Fig. S3)

### Microscopic investigations

Microscopic experiments were performed using an AxioImager microscope (Zeiss [Carl Zeiss], Thornwood, NY) coupled with a CoolSnap HQ camera (Roper Scientific) and a SpectraX LED lamp (Lumencor) at room temperature. Images were acquired and edited with MetaMorph (version 7.7.0.0; Universal Imaging). Strains were grown on glass slides covered with solid BMM and incubated for 12 to 24 hours. Co-localization of proteins was obtained by inoculating two different strains on the same BMM-coated glass slides in Petri dishes for 1 to 2 days. Hyphal fusion of both strains enabled the formation of heterokaryons by exchanging nuclei. GFP fluorescence and mRFP fluorescence were analyzed using filter sets (Chroma Technology Corp.) 49002 (GFP, excitation filter HQ470/40, emission filter HQ525/50, beamsplitter T495LPXR) and 49008 (mRFP, excitation filter HQ560/40, emission filter ET630/75m, beamsplitter T585lp). Septa in vegetative hyphae and ascogonial coils were stained using Calcofluor White M2R (CFW; Sigma [Sigma Chemical], St. Louis, MO) with a concentration of 1 μg/ml CFW stock solution diluted 1:400 in A. dest solution. CFW fluorescence was analyzed using Chroma filter set 31000v2 (excitation filter D350/50, emission filter D460/50, beam splitter 400dclp; Chroma Technology Corp., Bellows Falls, VT, USA).

To analyze the distribution of SmKIN3 on septa, 50 to 100 hyphal tips of the growing front were observed in 10 to 15 independent samples per strain in three different independent strains. Septal distances in hyphae were measured using MetaMorph (version 6.3.1; Universal Imaging). For analysis of hyper-septation, e.g. double and triple septa, we investigated at least 100 single hyphal branches for each strain. In the case of recombinant strains, three different single ascospore isolates were investigated to exclude side effects from random integration of recombinant DNA. In these cases, 600 hyphal branches were counted for each recombinant strain.

### Peptide selection for targeted quantification of phosphorylation sites

Based on the results of the global ITRAQ-based (phospho-)proteomic analyses previously performed (7,8), as well as unpublished work, phosphorylated peptides that showed differential regulation in STRIPAK mutant strains compared to wild type were selected and identified as putative dephosphorylation targets of the STRIPAK complex. In addition, the corresponding non-phosphorylated peptides were selected for quantification, thus enabling calculation of a site-occupancy value for the respective phosphorylation sites. As controls, phosphopeptides representing cell division control protein 48 (CDC48; SMAC_00109) and heat shock protein 90 (HSP90; SMAC_04445) were included (Figs. S25-S28), since they showed no regulation in the global experiments, and to our knowledge are not functionally connected to the STRIPAK complex.

### Synthesis, purification and quantification of SIS peptides

Synthesis of all SIS peptides was performed in-house using a Syro I synthesis unit (MultiSynTech, Witten, Germany) and Fmoc chemistry. Synthesis and subsequent purification were performed as described previously (42). Heavy-labeled lysine (^13^C_6_ ^15^N_2_) and arginine (^13^C_6_ ^15^N_4_) were incorporated at the C-terminus, and amino acid analysis was applied to determine peptide concentrations (43). A five-point calibration curve of derivatized amino acids, ranging from 5-25 pmol/μL, was used for quantification.

Protein extraction, digestion and normalization and phosphopeptide enrichment were performed as recently reported (7).

### nano-LC-MS/MS for PRM of unphosphorylated peptides

Samples were analyzed on an Ultimate 3000 nanroRSLC HPLC system coupled to a Q Exactive HF mass spectrometer (MS, both Thermo Scientific). The HPLC was equipped with a trapping column (100 μm x 2 cm C18, PepMap RSLC, Thermo Scientific) for pre-concentration, and an analytical column (75 μm x 50 cm C18, PepMap RSLC, Thermo Scientific) for separation of the peptides. Pre-concentration was performed for 5 min at a flow rate of 20 μL/min using 0.1% TFA, and separation was performed at a flow rate of 250 nL/min. An optimized binary gradient of solvent A (0.1% FA) and solvent B (84% acetonitrile, 0.1% FA) was used with the following steps: 0 min – 2%B; 5 min – 2%B; 10 min – 5%B; 50 min – 9%B; 73 min – 15%B; 100 min – 21%B; 115 min – 45%B, followed by two washing steps for 5 min at 95%B and 20 min of equilibration at 2%B. The MS was operated in PRM mode at a resolution of 60,000 (at 200 m/z) with a fixed first mass of 150 m/z. The AGC target was set to 1×10^6^ and a maximum injection time of 118 ms. Targeted precursors were isolated with a quadrupole isolation width of 0.4 m/z and fragmented with a normalized collision energy of 27. PRM acquisition was scheduled with a retention-time window of 2 min per target.

### nano-LC-MS/MS for PRM of phosphorylated peptides

Enriched phosphorylated peptides were analyzed with the same instrumentation and settings as described above. The LC gradient was optimized to suit the targeted phosphopeptides and the steps were modified as follows: 0 min – 3%B; 5 min – 3%B; 15 min – 7%B; 37 min – 10%B; 90 min – 20%B; 110 min – 27%B; 120 min – 45%B, followed by the same washing and equilibration steps as above.

### PRM development and analysis

To verify the linearity of response and determine the <u>l</u>imit <u>o</u>f <u>b</u>lank (LOB), LOD, and LLOQ response curves of both phosphorylated and non-phosphorylated SIS peptides were acquired. In both cases, a background matrix was generated by pooling aliquots of all individual samples. SIS peptides were spiked in at 8 different concentrations, covering a range of three orders of magnitude. Based on prior determination of individual SIS response factors, the highest concentrations for non-phosphorylated SIS peptides varied between 54 fmol and 2.73 pmol on-column, while the lowest concentrations were between 13 and 667 amol on-column. For the analysis of the individual samples, a total amount of non-phosphorylated SIS peptides ranging from 0.05 fmol to 60 fmol was spiked into 3 μg of total protein digest, depending on the expected endogenous concentration (all concentrations are given in Supplementary data file S1).

For phosphorylated SIS peptide calibration curve measurements, a total of 8 dilution points were generated with SIS peptide concentrations ranging from 600 amol to 2.4 pmol in 200 μg of background matrix for the best responding peptides, as detailed in supplementary data file S1.

In addition, six phoshphoisomeric <u>s</u>tandard <u>i</u>sotope-<u>l</u>abeled (SIL) peptides were included in the assay, but no endogenous peptide was detected. These peptides could be further used to rule out the presence of these isomers through knowledge of their retention time and diagnostic transitions. The median CV of the biological replicates was calculated as 16.3% for all non-phosphorylated and 16.1% for all phosphorylated peptides. The lowest average site occupancy was detected at 0.06% on S369 in the HSP 90 protein SMAC_04445. This site was included as negative control and its occupancy showed no significant difference between the wild type and any of the deletion strains. The phosphorylation site S2349 in the phosphatidylinositol 3-kinase TOR2 (SMAC_03322) exhibited the highest site occupancy with 92.2% measured in the wild type.

All calibration curve samples were subjected to TiO_2_-based phosphopeptide enrichment and nano-LC-MS/MS measurement in technical triplicates (16.7% of eluate per replicate) as detailed above. Additionally, two replicates without SIS spike-in (DS0) were processed in parallel to appropriately determine background signal levels. For the final analysis of individual samples, a total amount of 410 amol to 4.1 fmol was spiked into 450 μg of protein digest, followed by phosphopeptide enrichment and nano-LC-MS/MS measurement of 25% of the eluates after enrichment.

Skyline software (version 4.1, (44)) was used to analyze all of the PRM data. The top 3 most suitable transitions of every light and SIS peptide pair were chosen. All data were manually inspected for correct peak detection, retention time and integration, and peak areas were exported. R software (45) (version 3.5.3) was used for data analysis and calibration curve measurements, and LOB and LOD were calculated using the MSStats package (46). The L/H peak area ratios were used to determine the concentration (c) of phosphorylated and non-phosphorylated peptide per sample. The phosphorylation site occupancy was calculated using the following formula:

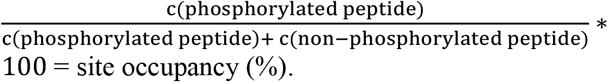

Statistical comparison of the phosphorylation site occupancy between wild type and knock-out strains was performed using a two-sided student’s t-test.

## Data availability

All targeted proteomics data and raw files are available through the Panorama repository (47) with the dataset identifier PXD023130 and via https://panoramaweb.org/SmKIN3.url

## Acknowledgements

We thank Ingeborg Godehardt and Susanne Schlewinski for superb technical help, and Varvara Solovyeva for help during her Bachelor thesis.

## Funding and additional information

VS receives a stipend from the Studienstiftung des Deutschen Volkes (German Academic Scholarship Foundation, Bonn Bad-Godesberg, Germany). This study was funded by the German Research Foundation (DFG) (Bonn Bad-Godesberg, Germany) (KU517/16-1, KU517/16-2, SI835/6-1, SI835/8-2).

## Abbreviations

BMM: biomalt-cornmeal medium
CFW: Calcofluor White
CR: cortical ring
CV: coefficient of variation
ELM: eukaryotic length motif
FHA: forkhead-associated
GCK: germinal centre kinase
iTRAQ: isobaric Tags for Relative and Absolute Quantitation
LFQ: label-free quantification
LLOQ: lower limit of quantification
LOB: limit of blank
LOD: limit of detection
NDR: nuclear DBF2-related
PRM: parallel reaction monitoring
SIL: standard isotope labeled
SIN: septation initiation network
SIP: SIN inhibitory PP2A
SIS: stable isotope-labeled standards
STRIPAK: striatin-interacting phosphatase and kinase
TMT: tandem mass tag

## References

1. Hwang, J., and Pallas, D. C. (2014) STRIPAK complexes: structure, biological function, and involvement in human diseases. Int J Biochem Cell Biol. 47, 118–148

2. Shi, Z., Jiao, S., and Zhou, Z. (2016) STRIPAK complexes in cell signaling and cancer. Oncogene. 35, 4549–4557

3. Gundogdu, R., and Hergovich, A. (2019) MOB (Mps one Binder) proteins in the Hippo pathway and cancer. Cells. 8

4. Kück, U., Radchenko, D., and Teichert, I. (2019) STRIPAK, a highly conserved signaling complex, controls multiple eukaryotic cellular and developmental processes and is linked with human diseases. Biol Chem. 400, 1005–1022

5. Roche, C. M., Loros, J. J., McCluskey, K., and Glass, N. L. (2014) *Neurospora crassa*: looking back and looking forward at a model microbe. Am J Bot. 101, 2022–2035

6. Hogrebe, A., von Stechow, L., Bekker-Jensen, D. B., Weinert, B. T., Kelstrup, C. D., and Olsen, J. V. (2018) Benchmarking common quantification strategies for large-scale phosphoproteomics. Nat Commun. 9, 1045

7. Märker, R., Blank-Landeshammer, B., Beier-Rosberger, A., Sickmann, A., and Kück, U. (2020) Phosphoproteomic analysis of STRIPAK mutants identifies a conserved serine phosphorylation site in PAK kinase CLA4 to be important in fungal sexual development and polarized growth. Mol Microbiol. 113, 1053–1069

8. Stein, V., Blank-Landeshammer, B., Müntjes, K., Märker, R., Teichert, I., Feldbrügge, M., Sickmann, A., and Kück, U. (2020) The STRIPAK signaling complex regulates dephosphorylation of GUL1, an RNA-binding protein that shuttles on endosomes. PLoS Genet. 16, e1008819

9. Marx, V. (2013) Targeted proteomics. Nat Methods. 10, 19–22

10. Wu, R., Dephoure, N., Haas, W., Huttlin, E. L., Zhai, B., Sowa, M. E., and Gygi, S. P. (2011) Correct interpretation of comprehensive phosphorylation dynamics requires normalization by protein expression changes. Mol Cell Proteomics. 10, M111 009654

11. Dekker, L. J. M., Zeneyedpour, L., Snoeijers, S., Joore, J., Leenstra, S., and Luider, T. M. (2018) Determination of site-specific phosphorylation ratios in proteins with targeted mass spectrometry. J Proteome Res. 17, 1654–1663

12. Prus, G., Hoegl, A., Weinert, B. T., and Choudhary, C. (2019) Analysis and interpretation of protein post-translational modification site stoichiometry. Trends in Biochemical Sciences. 44, 943–960

13. Radchenko, D., Teichert, I., Pöggeler, S., and Kück, U. (2018) A Hippo pathway-related GCK controls both sexual and vegetative developmental processes in the fungus *Sordaria macrospora*. Genetics. 210, 137–153

14. Simanis, V. (2015) *Pombe*’s thirteen - control of fission yeast cell division by the septation initiation network. J Cell Sci. 128, 1465–1474

15. Heilig, Y., Schmitt, K., and Seiler, S. (2013) Phospho-regulation of the *Neurospora crassa* septation initiation network. PLoS One. 8, e79464

16. Guertin, D. A., Chang, L., Irshad, F., Gould, K. L., and McCollum, D. (2000) The role of the sid1p kinase and cdc14p in regulating the onset of cytokinesis in fission yeast. EMBO J. 19, 1803–1815

17. Si, H., Rittenour, W. R., Xu, K., Nicksarlian, M., Calvo, A. M., and Harris, S. D. (2012) Morphogenetic and developmental functions of the *Aspergillus nidulans* homologues of the yeast bud site selection proteins Bud4 and Axl2. Mol Microbiol. 85, 252–270

18. Wu, H., Guo, J., Zhou, Y. T., and Gao, X. D. (2015) The anillin-related region of Bud4 is the major functional determinant for Bud4’s function in septin organization during bud growth and axial bud site selection in budding yeast. Eukaryot Cell. 14, 241–251

19. Gouw, M., Michael, S., Samano-Sanchez, H., Kumar, M., Zeke, A., Lang, B., Bely, B., Chemes, L. B., Davey, N. E., Deng, Z., Diella, F., Gurth, C. M., Huber, A. K., Kleinsorg, S., Schlegel, L. S., Palopoli, N., Roey, K. V., Altenberg, B., Remenyi, A., Dinkel, H., and Gibson, T. J. (2018) The eukaryotic linear motif resource - 2018 update. Nucleic Acids Res. 46, D428–D434

20. Sandrock, B., Böhmer, C., and Bölker, M. (2006) Dual function of the germinal centre kinase Don3 during mitosis and cytokinesis in *Ustilago maydis*. Mol Microbiol. 62, 655–666

21. Mayya, V., Rezual, K., Wu, L., Fong, M. B., and Han, D. K. (2006) Absolute quantification of multisite phosphorylation by selective reaction monitoring mass spectrometry: determination of inhibitory phosphorylation status of cyclin-dependent kinases. Mol Cell Proteomics. 5, 1146–1157

22. Schulze, W. X., Schneider, T., Starck, S., Martinoia, E., and Trentmann, O. (2012) Cold acclimation induces changes in *Arabidopsis* tonoplast protein abundance and activity and alters phosphorylation of tonoplast monosaccharide transporters. Plant J. 69, 529–541

23. Shi, T., Gao, Y., Gaffrey, M. J., Nicora, C. D., Fillmore, T. L., Chrisler, W. B., Gritsenko, M. A., Wu, C., He, J., Bloodsworth, K. J., Zhao, R., Camp, D. G., 2nd, Liu, T., Rodland, K. D., Smith, R. D., Wiley, H. S., and Qian, W. J. (2015) Sensitive targeted quantification of ERK phosphorylation dynamics and stoichiometry in human cells without affinity enrichment. Anal Chem. 87, 1103–1110

24. Aldous, S. H., Weise, S. E., Sharkey, T. D., Waldera-Lupa, D. M., Stuhler, K., Mallmann, J., Groth, G., Gowik, U., Westhoff, P., and Arsova, B. (2014) Evolution of the phosphoenolpyruvate carboxylase protein kinase family in C3 and C4 *Flaveria spp*. Plant Physiol. 165, 1076–1091

25. Yi, L., Shi, T., Gritsenko, M. A., X’Avia Chan, C. Y., Fillmore, T. L., Hess, B. M., Swensen, A. C., Liu, T., Smith, R. D., Wiley, H. S., and Qian, W. J. (2018) Targeted quantification of phosphorylation dynamics in the context of EGFR-MAPK pathway. Anal Chem. 90, 5256–5263

26. Chen, R., Xie, R., Meng, Z., Ma, S., and Guan, K. L. (2019) STRIPAK integrates upstream signals to initiate the Hippo kinase cascade. Nat Cell Biol. 21, 1565–1577

27. Heng, B. C., Zhang, X., Aubel, D., Bai, Y., Li, X., Wei, Y., Fussenegger, M., and Deng, X. (2020) An overview of signaling pathways regulating YAP/TAZ activity. Cell Mol Life Sci.

28. Bae, S. J., Ni, L., Osinski, A., Tomchick, D. R., Brautigam, C. A., and Luo, X. (2017) SAV1 promotes Hippo kinase activation through antagonizing the PP2A phosphatase STRIPAK. Elife. 6, e30278

29. Bae, S. J., Ni, L., and Luo, X. (2020) STK25 suppresses Hippo signaling by regulating SAV1-STRIPAK antagonism. Elife. 9

30. Heilig, Y., Dettmann, A., Mouriño-Pérez, R. R., Schmitt, K., Valerius, O., and Seiler, S. (2014) Proper actin ring formation and septum constriction requires coordinated regulation of SIN and MOR pathways through the germinal centre kinase MST-1. PLoS Genet. 10, e1004306

31. Singh, N. S., Shao, N., McLean, J. R., Sevugan, M., Ren, L., Chew, T. G., Bimbo, A., Sharma, R., Tang, X., Gould, K. L., and Balasubramanian, M. K. (2011) SIN-inhibitory phosphatase complex promotes Cdc11p dephosphorylation and propagates SIN asymmetry in fission yeast. Curr Biol. 21, 1968–1978

32. Justa-Schuch, D., Heilig, Y., Richthammer, C., and Seiler, S. (2010) Septum formation is regulated by the RHO4-specific exchange factors BUD3 and RGF3 and by the landmark protein BUD4 in *Neurospora crassa*. Mol Microbiol. 76, 220–235

33. Zhong, G., Wei, W., Guan, Q., Ma, Z., Wei, H., Xu, X., Zhang, S., and Lu, L. (2012) Phosphoribosyl pyrophosphate synthetase, as a suppressor of the sepH mutation in *Aspergillus nidulans*, is required for the proper timing of septation. Mol Microbiol. 86, 894–907

34. Bohnert, K. A., Grzegorzewska, A. P., Willet, A. H., Vander Kooi, C. W., Kovar, D. R., and Gould, K. L. (2013) SIN-dependent phosphoinhibition of formin multimerization controls fission yeast cytokinesis. Genes Dev. 27, 2164–2177

35. Westfall, P. J., and Momany, M. (2002) *Aspergillus nidulans* septin AspB plays pre- and postmitotic roles in septum, branch, and conidiophore development. Mol Biol Cell. 13, 110–118

36. Vargas-Muñiz, J. M., Renshaw, H., Richards, A. D., Waitt, G., Soderblom, E. J., Moseley, M. A., Asfaw, Y., Juvvadi, P. R., and Steinbach, W. J. (2016) Dephosphorylation of the core septin, AspB, in a protein phosphatase 2A-dependent manner impacts its localization and function in the fungal pathogen *Aspergillus fumigatus*. Front Microbiol. 7, 997

37. Mendoza, M., Redemann, S., and Brunner, D. (2005) The fission yeast MO25 protein functions in polar growth and cell separation. Eur J Cell Biol. 84, 915–926

38. Jerpseth, B., Greener, A., Short, J., Viola, J., and Kretz, P. (1992) XL1-blue MRF= *E. coli* cells: *mcrA-, mcrCB-, mcrF-, mmr-, hsdR-* derivative of XL1-blue cells. Mol Biol 5, 81–83

39. Sambrook, J., and Russel, D. (2001) Molecular cloning: a laboratory manual, Cold Spring Harbor Laboratory Press, NY

40. Engh, I., Würtz, C., Witzel-Schlömp, K., Zhang, H. Y., Hoff, B., Nowrousian, M., Rottensteiner, H., and Kück, U. (2007) The WW domain protein PRO40 is required for fungal fertility and associates with woronin bodies. Eukaryot Cell. 6, 831–843

41. Dirschnabel, D. E., Nowrousian, M., Cano-Dominguez, N., Aguirre, J., Teichert, I., and Kück, U. (2014) New insights into the roles of NADPH oxidases in sexual development and ascospore germination in *Sordaria macrospora*. Genetics. 196, 729–744

42. Dickhut, C., Feldmann, I., Lambert, J., and Zahedi, R. P. (2014) Impact of digestion conditions on phosphoproteomics. J Proteome Res. 13, 2761–2770

43. Cohen, S. A., and Michaud, D. P. (1993) Synthesis of a fluorescent derivatizing reagent, 6-aminoquinolyl-N-hydroxysuccinimidyl carbamate, and its application for the analysis of hydrolysate amino acids via high-performance liquid chromatography. Anal Biochem. 211, 279–287

44. MacLean, B., Tomazela, D. M., Shulman, N., Chambers, M., Finney, G. L., Frewen, B., Kern, R., Tabb, D. L., Liebler, D. C., and MacCoss, M. J. (2010) Skyline: an open source document editor for creating and analyzing targeted proteomics experiments. Bioinformatics. 26, 966–968

45. Team, R. C. (2016) R: A Language and Environment for Statistical Computing. in R Foundation for Statistical Computing, Vienna, Austria.

46. Choi, M., Chang, C. Y., Clough, T., Broudy, D., Killeen, T., MacLean, B., and Vitek, O. (2014) MSstats: an R package for statistical analysis of quantitative mass spectrometry-based proteomic experiments. Bioinformatics. 30, 2524–2526

47. Sharma, V., Eckels, J., Schilling, B., Ludwig, C., Jaffe, J. D., MacCoss, M. J., and MacLean, B. (2018) Panorama public: A public repository for quantitative data sets processed in skyline. Mol Cell Proteomics. 17, 1239–1244

